# Which actin genes are necessary for zebrafish heart development and function?

**DOI:** 10.1101/2020.08.25.266932

**Authors:** Kendal Prill, Matiyo Ojehomon, Love Sandhu, Suchandrima Dutta, John F. Dawson

## Abstract

Heart failure is the number one cause of mortality in the world, contributed to by cardiovascular disease. Many diseases of the heart muscle are caused by mutations in genes encoding contractile proteins, including cardiac actin mutations. Zebrafish are an advantageous system for modeling cardiac disease since embryos can develop without a functional heart. However, genome duplication in the teleost lineage creates a unique obstacle by increasing the number of genes involved in heart development. Four actin genes are expressed in the zebrafish heart: *acta1b; actc1c;* and duplicates of *actc1a* on chromosome 19 and 20. Here, we characterize the actin genes involved in early zebrafish heart development using *in situ* hybridization and CRISPR targeting to determine which gene is best to model changes seen in human patients with heart disease. The *actc1a* and *acta1b* genes are predominant during embryonic heart development, resulting in severe cardiac phenotypes when targeted with CRISPRs. Targeting these two cardiac genes with CRISPRs simultaneously results in a more severe phenotype than their individual counterparts, with the results suggesting compensation for lost actin genes by other actin paralogues. Given the duplication of the *actc1a* gene, we recommend *acta1b* as the best gene for targeted cardiac actin research.

## Introduction

Heart failure is the number one cause of death worldwide, with cardiovascular disease being a major contributor^1^. Cardiomyopathies are called “diseases of the sarcomere” because mutations in genes encoding the proteins of the sarcomere contractile machinery are a main cause of cardiomyopathies, including myosin, troponin, tropomyosin, cardiac myosin binding protein C, and cardiac actin (ACTC)^2,3^. Recent efforts have targeted some of these proteins for drug development to treat cardiomyopathies^4^.

Testing drugs with a whole animal system is vital for developing treatments for diseases. We seek to understand how changes in the cardiac actin gene (*ACTC*) in people lead to different cardiomyopathies. We have studied several ACTC variants at the molecular level^5–9^; however, our goal is to integrate our molecular knowledge of ACTC biochemical changes with physiological dysfunction in a whole animal by gene editing the cardiac actin gene in the model organism.

The zebrafish is an excellent model for cardiac research^10,11^: their embryos are transparent with heartbeats detectable at 24 hours post-fertilization (hpf), and they do not require a fully functional heart for viability for the first 5 days post-fertilization (dpf) due to diffusion of oxygen through the tissues; hence, embryonic lethal heart mutations seen in mammals can be studied more readily in zebrafish. In addition, zebrafish are small and easy to maintain, with quick growth, large numbers of progeny, and the genome of the Tübingen (Tü) strain has been completely sequenced.

However, unlike the single a-cardiac actin gene (*ACTC*) in humans, the zebrafish genome contains four actin genes that are expressed in the heart *(zfactc* genes): *acta1b; actc1c;* and duplicates of *actcla* on chromosome 19 and 20. Cardiac-related effects of mutations in *actc1a* and *acta1b* have been studied previously^12–14^, while we identified and performed preliminary characterization of *actc1c*^15^.

Shih et al. (2015) performed transcriptome analysis of *actc1a* and *acta1b* in embryonic (96 hpf) and adult (6 month-old) zebrafish hearts, showing higher *acta1b* expression in early development while *actc1a* is expressed at higher levels during adulthood, suggesting developmental regulation of *zfactc* genes^16^. Functional and spatiotemporal characterization of these *zfactc* genes is needed to assign a cardiac designation with high confidence.

While early expression of *actc1a* and *acta1b* has been characterized with *in situ* hybridization (ISH), this previous work focused primarily on the somites of the tail. The *actc1a* gene was the subject of expression characterization in the somites and included the heart at the 1-4 somites to 7 dpf stages^16–18^, with *acta1b* ISH analysis focused on the somites^18–20^. Our preliminary characterization of *actc1c* showed expression in the heart and the somites at 36 hpf^15^. Owing to teleost gene duplication, duplicate *actc1a* genes located on chromosomes 19 and 20 are identical in sequence up to about 600 bp before the start codon, making designing *in situ* hybridization probes or CRISPR sgRNA that distinguish between the duplicate genes extremely challenging.

To determine the *zfactc* genes necessary for zebrafish heart development and function and which is the best to edit and model human cardiac diseases resulting from *ACTC* mutations, we studied the spatiotemporal expression of these genes in the heart, employing *in situ* hybridization in embryos from 24 hpf to 96 hpf and observing the functional consequences of CRISPRs targeting the genes. All three *zfactc* genes are expressed in the heart at initial stages of heart development; however, the *actc1a* and *acta1b* genes are the predominant paralogues expressed during embryonic heart development and result in severe cardiac phenotypes when targeted with CRISPRs. The *actc1c* gene seems to be a minor player, with its expression occurring primarily in the first 2 dpf. At the same time, CRISPR work targeting *actc1c* results in a cardiac phenotype, suggesting that this gene plays a role in cardiac development. Targeting two *zfactc* genes with CRISPRs simultaneously results in a more severe phenotype than their individual counterparts, with the results suggesting compensation for lost *zfactc* genes by other actin paralogues. Given the gene duplication of the *actc1a* gene, we suggest that the *acta1b* gene is the best candidate for cardiac actin research.

## Materials and Methods

### Ethics Statement

All protocols were carried out according the guidelines stipulated by the Canadian Council for Animal Care and the University of Guelph’s Office of Research Animal Care Committee (Animal Use Protocol license: 4309)

### Zebrafish Maintenance

Adult zebrafish (Tübingen strain) were maintained according to guidelines by the Canadian Council on Animal Care and kept on a 12/12-hour light and dark cycle at 28°C. Adults were fed brine shrimp (Hikari Bio-Pure Brine Shrimp) and fish flakes (Omega One) daily in a cycled-water aquatic facility. Embryos were collected from crossing wild-type adult zebrafish and grown at 28°C in zebrafish embryo medium^21^ for up to 6 days prior to fixation.

### In situ hybridization

Zebrafish embryos were staged and fixed in 4% paraformaldehyde/PBS overnight at 4°C. Cardiac actin probes were cloned into TOPO-plasmids (Thermofisher) for probe synthesis (Table 1). Antisense RNA probes were synthesized from the TOPO construct using SP6 RNA polymerase (Thermofisher).

**Table 1:**
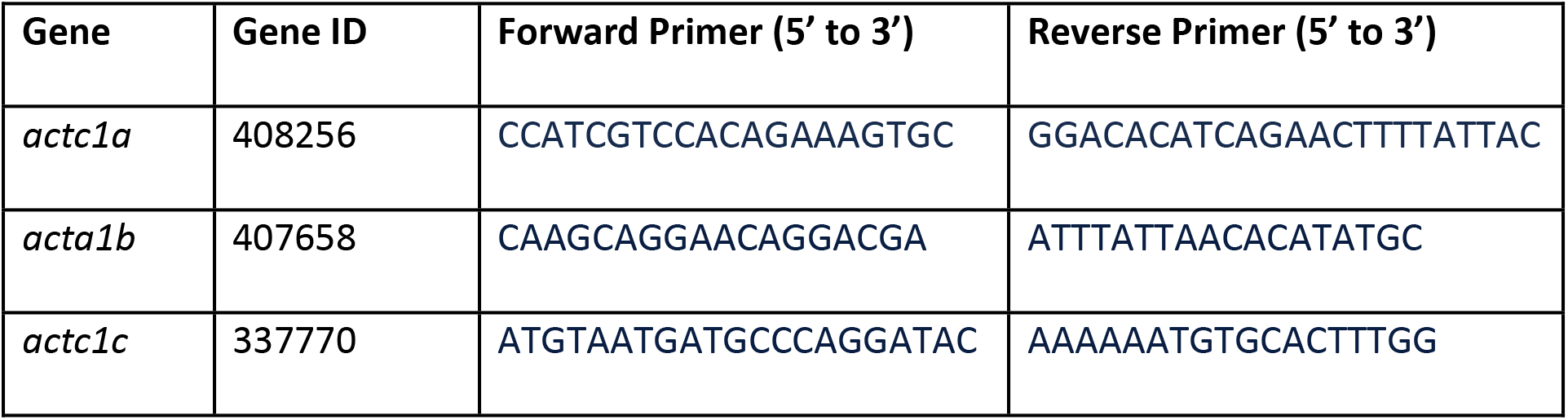
Primers used to amplify *in situ* hybridization probes.

*In situ* hybridizations were carried out as previously described^22^ using an Intavis *In Situ* Pro Liquid Handling Robot (Intavis, Koeln) with the exception of the proteinase K digestion and staining reaction steps, which were performed by hand.

### CRISPR sgRNA Preparation and Microinjections

CRISPR single guide RNA (sgRNA) was designed for *actc1a, actc1c* and *acta1b* using CHOPCHOP (https://chopchop.cbu.uib.no; danRer11/GRCz11)^23^, selecting the sgRNA that returned the fewest off-target sites (Table 2). Given the extreme identity between the two *actc1a* genes physically located on chromosomes 19 and 20, one sgRNA was designed that targets both genes.

**Table 2:**
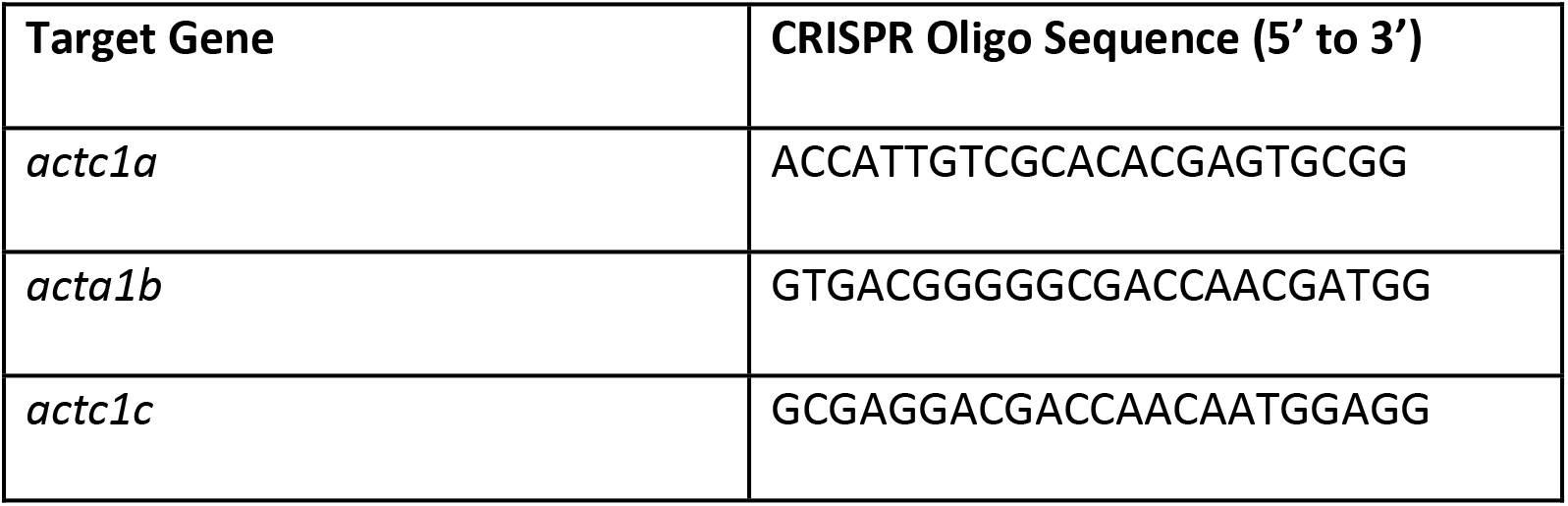
CRISPR single guide RNA designed to target the *zfactc* genes.

sgRNA was synthesized using SP6 RNA Polymerase (Thermofisher). Cas9 (1 mg/ml; CP01-200, PNA Bio Inc) and fresh sgRNA (1 ug) was injected into 1-cell zebrafish embryos that were allowed to recover in zebrafish embryo medium at 28°C. Zebrafish embryos were monitored daily for the appearance of phenotypes and imaged using an iPhone 6 camera (8 megapixel, 1080p HD video at 60 fps; Apple Inc.)

### High Resolution Melt Curve Analysis Screening and Sequencing

For screening using High Resolution Melt Curves (HRM), genomic DNA was extracted from zebrafish embryos by individually lysing tissue in 10 ul of 0.5 M NaOH at 95°C for 45 mins, followed by neutralization with 0.2 mM Tris-HCl (pH 8). DNA was diluted 1/20 for best amplification results during HRM. The HRM amplicons were designed to be no larger than 200 bp and centered on the CRISPR-Cas9 cut site (Table 3). High Resolution Melt Curves were produced using the saturating dye, EvaGreen (Type-it HRM PCR Kit, Qiagen) and the manufacturers recommended protocol for use. HRM was performed using a StepOne Plus Thermocycler (Thermofisher) and results interpreted using High Resolution Melt Software v3.0.1 (Thermofisher).

**Table 3:**
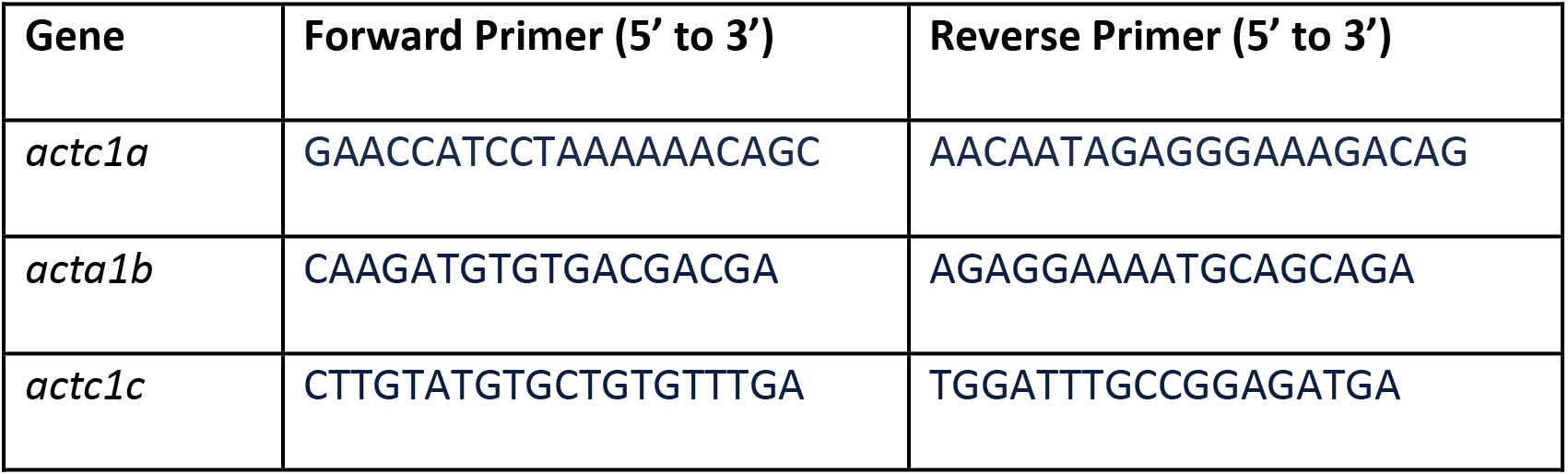
Primers employed for high resolution melting analysis of *zfactc* amplicons.

Embryos that demonstrated melt curves with melting temperature at 50% of the maximum temperature (Melt50) values that differed at least 0.5°C from their un-injected control siblings were sent for sequencing. Positive HRM samples were isolated using a PCR purification kit (Qiagen) and submitted to the University of Guelph, Agriculture and Food Laboratory for Sanger Sequencing. Sequences were analyzed using TIDE^24^ and Gear-Indigo (https://www.gear-genomics.com) to identify nucleotide changes between un-injected CRISPR control and CRISPR-injected samples.

### Heart Rate Acquisition and Analysis

The heart rates of all CRISPR-injected embryos were video-recorded daily from 1-6 dpf using an iPhone 6 camera (8 megapixel, 1080p HD video at 60 fps; Apple Inc.). Captured videos were analyzed with DanioScope software (Noldus) to determine heart rates. Genomic DNA was extracted from these embryos and screened for mutations using HRM as above and hits were sent for Sanger Sequencing. Embryos with mutated sequences had their heart rates graphed in comparison to wild-type controls.

## Results

### actc1a and acta1b are the predominant actin paralogues expressed during striated muscle development

To identify which actin paralogues are necessary for heart development, we analyzed the expression of *actc1a, acta1b* and *actc1c* with *in situ* hybridization (ISH) during early stages of embryogenesis (Fig 1). At 24 hpf, all 3 actin genes are expressed in the linear heart tube (Fig 1, A&B; white arrowheads), although *actc1c* appears to be limited to ventricle cardiomyocytes (Fig 1C)^25^. *actc1a* is expressed in the zebrafish heart at all stages examined with the strongest expression observed in the ventricle (Fig 1, A,D,E,J,K,P&O). The continued expression of *actc1a* throughout embryogenesis suggests *actc1a* is required for cardiac muscle development.

**Figure 1.**
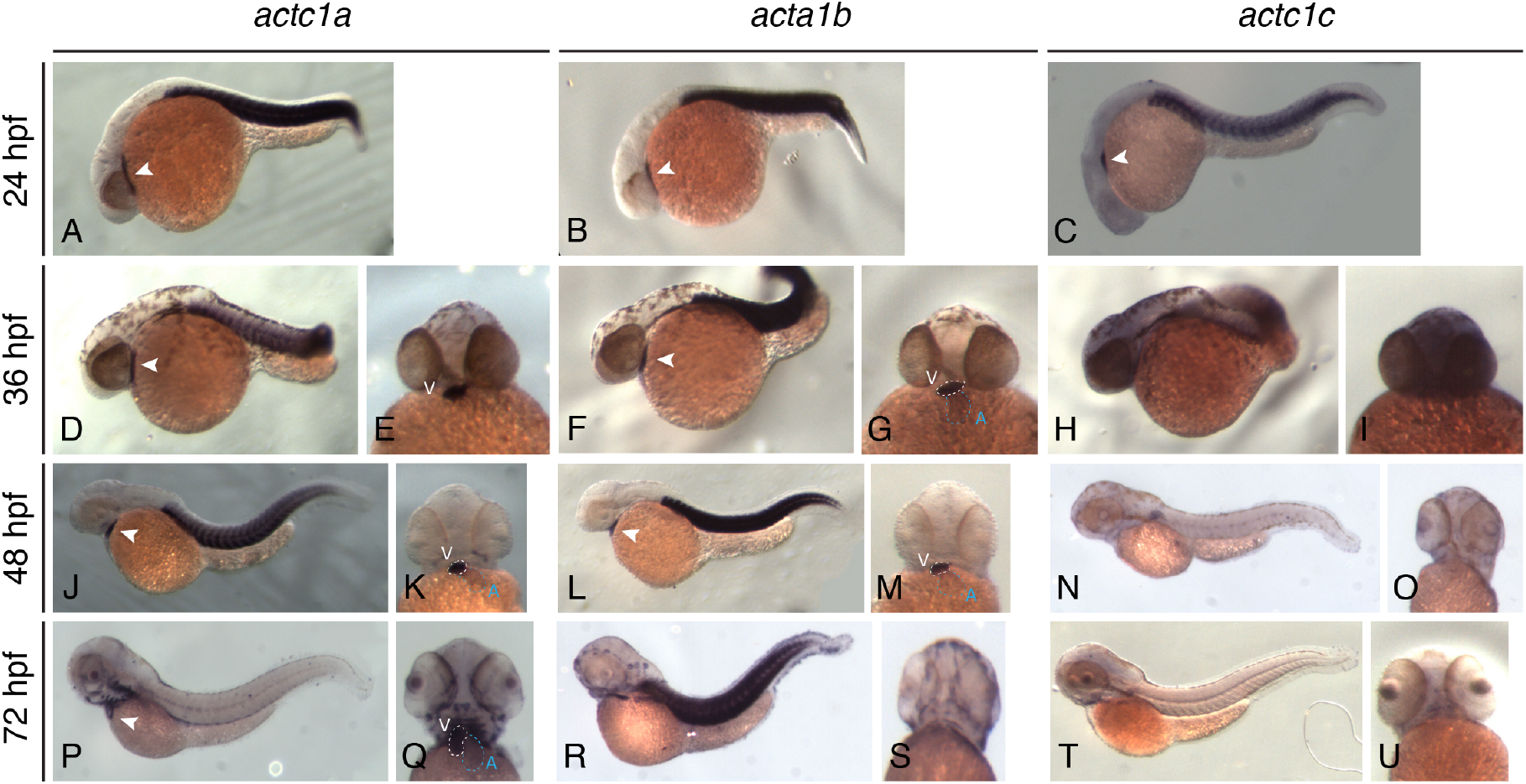
Expression of three actin paralogues throughout early embryo development. At 24 hpf, *in situ* hybridization reveals somite and heart tube-restricted expression of *actc1a* (A), *acta1b* (B) and *actc1c* (C). By 36 hpf, *actc1a* becomes restricted further to the ventricle (D, E) while *acta1b* is mainly expressed in the ventricle and faintly in the atrium (F, G). *Actc1C* demonstrates a ubiquitous expression, with the exception of the heart, by 36 hpf (H, I). At 48 hpf, *actc1a* (J, K) and *acta1b* (L, M) share nearly identical expression in the heart and somites with the atrium displaying low expression for both paralogues. By 48 hpf, *actc1c* expression is only observed in the pectoral fin buds (N, O). *Actcla* expression becomes restricted to the heart and head muscles at 72 hpf (P, Q). At 72 hpf, *acta1b* is only expressed in the head muscles and somites with no observable expression in the heart (R, S). By 72 hpf, *actc1c* is not expressed in the striated muscle of the developing zebrafish embryo (T, U). (white arrowheads = heart expression; v = ventricle, a = atrium; white dotted lines outline ventricle; blue dotted lines outline atrium).

Similar to *actc1a*, *acta1b* is also expressed in the heart for significant stages of heart development. *acta1b* is expressed strongly in the ventricle and weakly in the atrium from 24-48 hpf (Fig 1, B,F,G,L&M). By 72 hpf, *acta1b* is no longer expressed in the heart and is restricted to the skeletal muscle of the head and trunk (Fig 1, R&S). Unlike *actc1a, acta1b* is only expressed during the stages of heart morphogenesis and cardiac sarcomere formation, suggesting *acta1b* may be required for the assembly but not the maintenance of the zebrafish heart muscle.

*actc1c* expression is not observed in the heart at stages beyond 24 hpf (Fig 1, C,H&I) and ceases expression in skeletal muscle by 48 hpf (Fig 1, N,O,T&U). This lack of cardiac expression after 24 hpf suggests that *actc1c* is not required throughout heart development but does not rule out the possibility that *actc1c* may be required for the initial formation of cardiac sarcomeres in the ventricle.

### actc1a, acta1b and actc1c are all required for normal heart development and function

We demonstrated that *actc1a, acta1b* and *actc1c* are expressed at the earliest stages of heart development at 24 hpf but have different temporal and spatial expression patterns from 36-72 hpf (Fig 1). Since all three paralogues were expressed in the developing linear heart tube, we tested the necessity for each actin gene in heart development by disrupting each paralogue using the CRISPR/Cas9 system.

All CRISPR/Cas9-injected embryos displayed similar cardiac phenotypes ranging from normal to significantly reduced heart rates, and unlooped and minor to severely degenerated hearts. Embryos injected with either *acta1b-* or *actc1c*-targeting CRISPR sgRNA displayed pericardial edema when compared to mock-injected and embryos injected with *actc1a*-targeting CRISPR sgRNA embryos at 72 hpf (Fig 2, A-D; black arrowheads). At 96 hpf, *actc1a*-CRISPR-injected embryos demonstrated pericardial edema like *acta1b* or *actc1c*-injected embryos (Fig 2, C-F).

**Figure 2.**
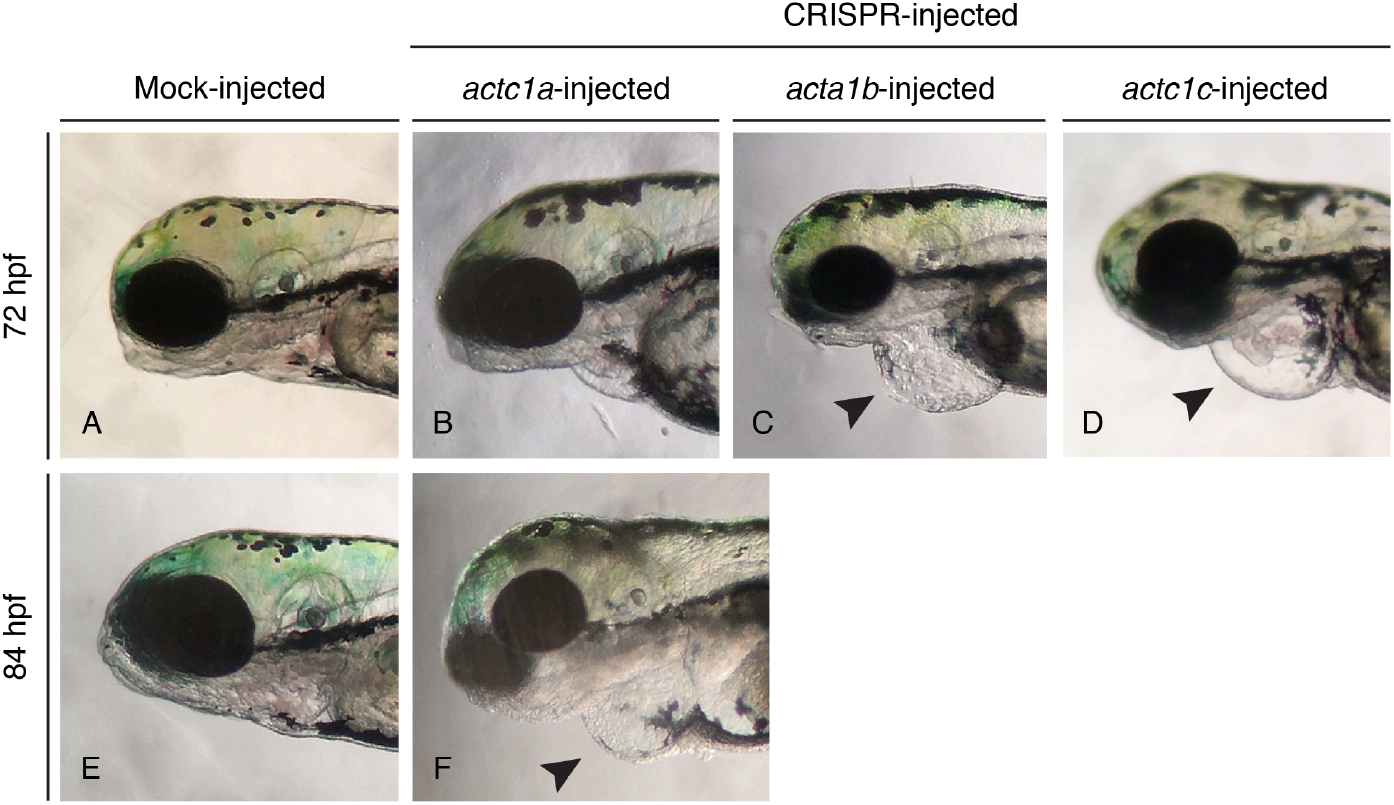
Phenotypes of cardiac actin CRISPR-injected embryos. When compared to mock-injected control embryos (A), embryos injected with *actc1a-*targeting CRISPR sgRNA (B) displayed only a slight pericardial edema while embryos injected with *acta1b*-targeting (C) or *actc1c-* targeting CRISPR sgRNA (D) demonstrated pericardial edema (black arrowheads) and incompletely looped hearts. By 96 hpf, embryos injected with *actc1a-*targeting CRISPR sgRNA exhibited pericardial (black arrowhead) and yolk sac edema when compared to mock-injected control embryos (E, F).

Although the severity of cardiac phenotypes varied across embryos injected with the same CRISPR sgRNA, we suspect genetic mosaicism produced by CRISPR/Cas9 cutting and non-homologous end joining repair accounts for this variation.

### Simultaneous CRISPR/Cas9 against actc1a and acta1b results in more severe cardiac phenotypes

We have shown that *actc1a* is expressed at all early stages of heart development when compared to either *acta1b* or *actc1c,* yet a *actc1a*-CRISPR-mutant phenotype appears later than the cardiac phenotypes of *acta1b* or *actc1c* (Fig 2). Based on these data, we hypothesize that *acta1b* and possibly *actc1c,* compensates for the absence of *actc1a* protein in initial stages of development during *acta1b* expression up to 72 hpf. To test this hypothesis, we injected embryos with both *actc1a-* and *acta1b-* targeting CRISPR sgRNA and compared their heart rates to wild-type and embryos injected with either *actc1a-* or *acta1b*-targeting CRISPR sgRNA alone. We would expect double *actc1a-acta1b* mutants to display heart rates similar to *acta1b*-CRISPR mutant heart rates at 72 hpf since expression of both predominant cardiac actin genes is absent and *acta1b* cannot compensate for *actc1a.*

At 48 and 72 hpf, the double *actc1a*-*acta1b*-targeting CRISPR sgRNA-injected embryos did not display significantly different heart rates when compared to wild-type, *actc1a-* or *acta1b*- targeting CRISPR sgRNA-injected embryos (Fig 3). However, by 96 hpf, the average heart rates of *actc1a-acta1b*-targeting CRISPR sgRNA-injected embryos were significantly lower than the heart rates observed with wild-type embryos. The difference between wild-type and *zfactc*-targeting CRISPR sgRNA-injected embryo heart rates continues to increase as embryogenesis progresses, suggesting that the hearts of embryos injected with *zfactc*-targeting CRISPR sgRNA do not continue proper development.

**Figure 3.**
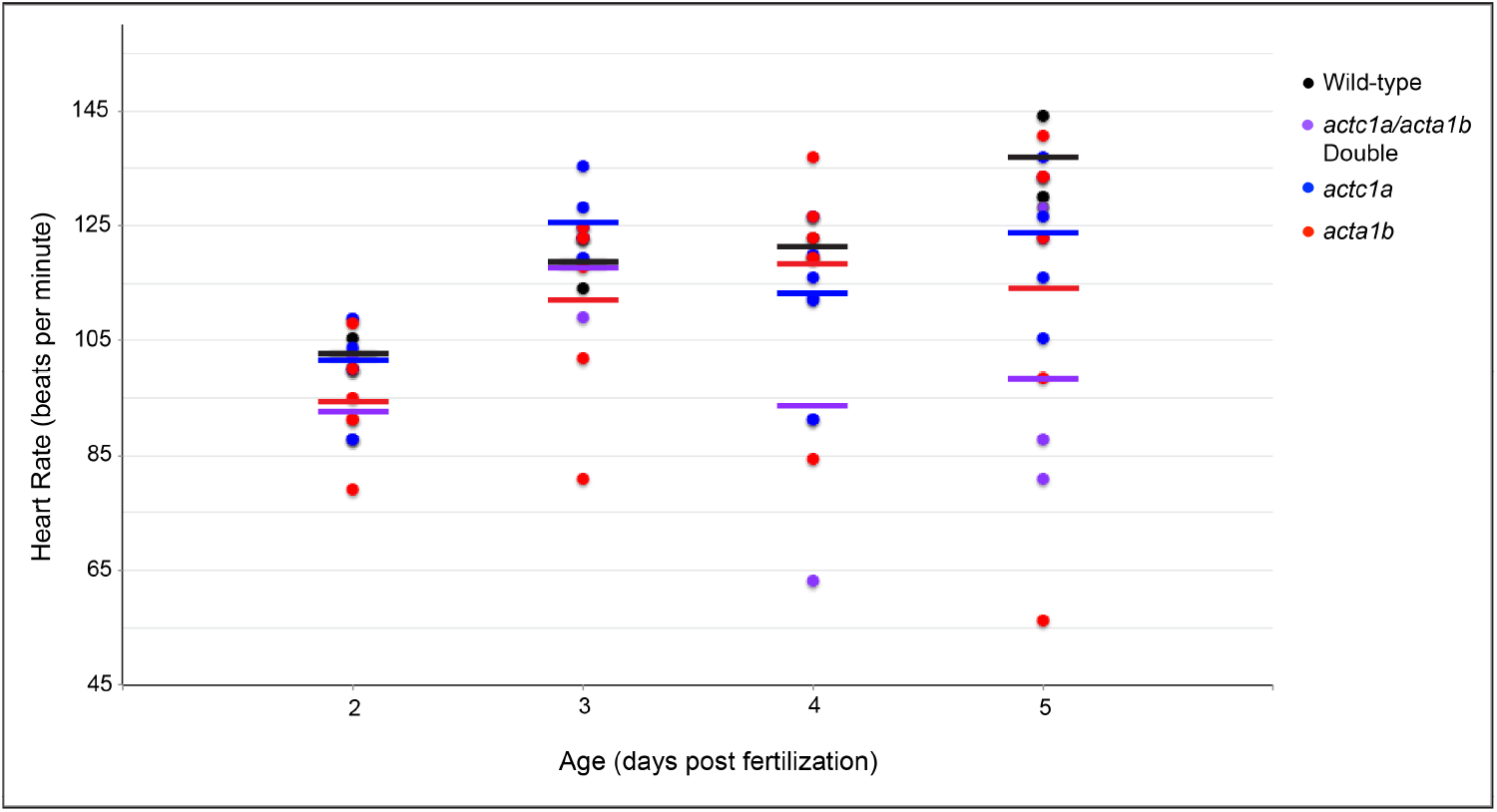
Heart Rates of embryos injected with cardiac actin-targeting CRISPR sgRNA throughout embryogenesis. The heart rates of wild-type and genotype-confirmed embryos injected with cardiac actin-targeting CRISPR sgRNA were recorded daily from 2-5 days. Individual (circles) and average (lines) heart rates are displayed for each genotype at every age. With the exception of 72 hpf, embryos injected with cardiac actin-targeting CRISPR sgRNA demonstrate a lower average heart rate when compared to control wild-type embryos. By 96 hpf, the heart rate of the double *actc1a-acta1b-targeting* CRISPR sgRNA-injected embryos have significantly lower heart rates than embryos injected with either *actc1a-* or *acta1b-*targeting CRISPR sgRNA. (genotype confirmed: wild-type, n=2; *actc1a/acta1b,* n=3; *actc1a,* n=6, *acta1b,* n=6).

## Discussion

Zebrafish are an excellent system for modeling human cardiac disease and dissecting the mechanisms behind disease progression and treatment. However, a genome duplication event unique to the teleost lineage can complicate specific gene targeting studies due to genetic compensation. Zebrafish cardiac actin genes are no exception to this complication with 4 identified genes (*actc1a* (chromosome 19 and 20), *acta1b* and *actc1c*) contributing to heart development. Based on *in situ* expression (Fig 1), and previous data^16^, *actc1a* has continuous transcription in the heart throughout development; however, *actc1a* is present as two identical genes on chromosomes 19 and 20. Sequencing of these regions reveals extreme sequence identity between the two occurrences of *actc1a,* so modifying and analyzing one *actc1a* isoform is very challenging without modifying one isoform first to differentiate the two.

*acta1b* is expressed in both chambers during the initial stages of heart development that include formation of the cardiac sarcomeres and heart looping. Additionally, there is only one *acta1b* gene, making targeting and analysis of human cardiac mutations more efficient and feasible than *actc1a* (19/20). *actc1c* demonstrated a very brief expression profile in the zebrafish heart, suggesting it is not as necessary for heart development as the other *zfactc* genes.

When we combine the expression profiles with the phenotypes of CRISPR-targeted *zfactc* genes, *acta1b* is required for normal heart development. When *acta1b* was targeted by CRISPR, cardiac phenotypes manifested early and became progressively worse over time (Fig 2 and 3). Targeting *actc1a* (19/20) with CRISPR resulted in phenotypes that appeared later than *acta1b* but followed a similar phenotype progression. The phenotypic delay with *actc1a* targeting suggests that *acta1b* and/or *actc1c* compensate for the lack of functional *actc1a.* We also considered that one wild-type isoform of *actc1c* (19 or 20) could still be expressed in these CRISPR mutants. However, due to the identical sequences of the two *actc1a* genes, we would hypothesize that we achieved a minimum threshold for completely modifying both *actc1a* genes within CRISPR-injected embryos. Simultaneously targeting *acta1b* and *actc1a* with CRISPR/Cas9 resulted in significantly worse heart phenotypes than targeting either single gene (Fig 3), supporting a compensation model by cardiac actin paralogues.

Taken together, our data suggests *acta1b* is the best zebrafish cardiac actin gene for modeling human heart diseases resulting from mutations in the cardiac actin gene. *acta1b* mutants demonstrated minimal compensation by other *zfactc* gene early in embryogenesis; ideal conditions for characterizing the phenotype and disease mechanism of a specific human actin mutation.

This work provides a foundation to model human actin mutations in zebrafish. Future work will focus on introducing human cardiac actin mutations into *acta1b* and characterizing the disease as well as exploring methods of treatment. Additionally, determining the changes in cardiac actin paralogue expression in response to single actin knockouts would further dissect the actin gene compensation hypothesis.

## Acknowledgements

This work was funded by a Heart and Stroke Foundation of Canada Grant-in-Aid to JFD (G-18-0020424).

